# RNA:DNA hybrids in the human genome have distinctive nucleotide characteristics, chromatin composition, and transcriptional relationships

**DOI:** 10.1101/020545

**Authors:** Julie Nadel, Rodoniki Athanasiadou, Christophe Lemetre, N. Ari Wijetunga, Pilib Ó Broin, Hanae Sato, Zhengdong Zhang, Jeffrey Jeddeloh, Cristina Montagna, Aaron Golden, Cathal Seoighe, John M. Greally

## Abstract

**Background:** RNA:DNA hybrids represent a non-canonical nucleic acid structure that has been associated with a range of human diseases and potential transcriptional regulatory functions. Mapping of RNA:DNA hybrids in human cells reveals them to have a number of characteristics that give insights into their functions.

**Results:** We find RNA:DNA hybrids to occupy millions of base pairs in the human genome. A directional sequencing approach shows the RNA component of the RNA:DNA hybrid to be purine-rich, indicating a thermodynamic contribution to their *in vivo* stability. The RNA:DNA hybrids are enriched at loci with decreased DNA methylation and increased DNase hypersensitivity, and within larger domains with characteristics of heterochromatin formation, indicating potential transcriptional regulatory properties. Mass spectrometry studies of chromatin at RNA:DNA hybrids shows the presence of the ILF2 and ILF3 transcription factors, supporting a model of certain transcription factors binding preferentially to the RNA:DNA conformation.

**Conclusions:** Overall, there is little to indicate a dependence for RNA:DNA hybrids forming co-transcriptionally, with results from the ribosomal DNA repeat unit instead supporting the intriguing model of RNA generating these structures *in trans.* The results of the study indicate heterogeneous functions of these genomic elements and new insights into their formation and stability *in vivo.*

## BACKGROUND

The complex regulatory process leading to gene expression involves, as a major upstream influence, the effects of transcription factors (TFs) binding to specific DNA motifs. The targeting of TFs to specific locations is an informational puzzle, as the number of potential binding sites represented by their generally short sequence binding motifs vastly exceeds the minority used *in vivo.* This observation suggests that there is additional information present in genomic organization that determines the selection of this subset of sequence motifs. Studies aiming to identify these extra layers of genomic information have revealed influences of chromatin organization [3-6] and DNA methylation [6-8], each of which can facilitate or reduce TF binding to cognate motifs, but the role of the conformation of the DNA molecule *in vivo* is less well studied. While it is known that nucleic acids can form numerous non-canonical conformations [9], the influence of these conformations in living cells remains under-studied. There is, however, evidence from *in vitro* assays that DNA conformation influences binding of proteins [10]. As examples, the SP1 transcription factor binds preferentially to the intra-strand G-quadruplex structure *in vitro* [11], while we have found the methyl-binding domain of the Mecp2 protein to bind preferentially to single-stranded DNA (ssDNA), also *in vitro* [12]. These observations indicate that the exploration of these and other non-canonical structures occurring *in vivo* may be fruitful in adding a layer of information to the transcriptional regulatory processes. The potential for ssDNA to occur in living cells, prompted by the results of our Mecp2 studies [12], raised the question about how such structures could be created and maintained stably *in vivo.* One candidate process to mediate the stable formation of ssDNA is the generation of an RNA:DNA hybrid on one DNA strand leaving the other strand in a single-stranded conformation, a nucleic acid structure referred to as an R-loop [13].

Formation of an R-loop has multiple potential consequences in terms of local organization of transcriptional regulatory elements. The helical conformation of the RNA:DNA hybrid differs from the B-form typical of double-stranded DNA (dsDNA), instead creating a conformation intermediate with the A-form associated with dsRNA [14]. A locus forming an RNA:DNA hybrid therefore creates a double-stranded A/B intermediate conformation, with a second target for single-stranded nucleic acid binding proteins on the complementary, displaced DNA strand. Another property of the R-loop is the displacement by the RNA of G-rich ssDNA [15, 16], allowing the formation of intramolecular G-quadruplex structures [17]. The potential that RNA:DNA hybrids may be resistant to the activity of DNA methyltransferases has previously been proposed [18], as has their failure to organize DNA into a nucleosomal conformation [19], further adding to their local influence on nucleic acid organization.

Formation and maintenance of an RNA:DNA hybrid is subject to many influences [15, 16, 20-23]. Transcription of a locus has been positively associated with RNA:DNA hybrid formation [24, 25], presumably by the RNA acting *in cis* with the DNA from which it was transcribed, but there is evidence in yeast that Rad51 can facilitate RNA molecules *in trans* also forming RNA:DNA hybrids [26]. Mutations of enzymes such as RNase H [27], which specifically hydrolyzes the RNA in RNA:DNA hybrids, RNA helicases [28] and topoisomerases [29] have been found to be associated with the increased formation of RNA:DNA hybrids, supporting a model in which these enzymes normally function to remove these structures from the genome. The presence of RNA:DNA hybrids at ribosomal DNA repeats appears to be a conserved feature from yeast [30] to human cells [18], for which any associated physiological role remains unclear. Functionally, RNA:DNA hybrids and their associated ssDNA regions have been found to have numerous properties *in vitro* and *in vivo* in a range of organisms. These include involvement in immunoglobulin class switching [31, 32], regulation of gene expression [33], constitutive formation in yeast telomeres [34] and at the origin of replication in mitochondrial DNA [23]. Additionally, these structures have been linked with epigenetic modifications, such as chromatin organization through enrichment at condensed chromatin marked by histone H3 serine 10 (H3S10) phosphorylation in yeast, *C. elegans* and human HeLa cells [35], centromeric heterochromatin [36], and formation at promoter CpG islands lacking DNA methylation [18]. The functions attributed to RNA:DNA hybrids are thus diverse and appear to have a major degree of dependence upon their genomic context.

RNA:DNA hybrids are being increasingly associated with human diseases, with a major concern that their presence predisposes a locus to chromosomal breakage. For example, it has been shown that R-loops are processed by the nucleotide excision repair endonucleases XPF and XPG into double strand breaks [37], and both BRCA1 [38] and BRCA2 [39] have been implicated as major processing enzymes involved in the resolution of RNA:DNA hybrids. The formation of RNA:DNA hybrids has also been associated with a number of neurological diseases. Mutations in the RNA:DNA helicase senataxin (*SETX*) mutations are implicated in the dominant juvenile form of amyotrophic lateral sclerosis type 4 (ALS4) and a recessive form of ataxia oculomotor apraxia type 2 (AOA2) [40]. RNase H2 (*RNASEH2*) mutations are among those associated with Aicardi-Goutières syndrome, in which the accumulation of unusual nucleic acids triggers inflammatory and autoimmune responses [41]. In addition, it is known that triplet repeats are prone to forming unusual nucleic acid structures, including R-loops and RNA:DNA hybrids, a phenomenon conserved in organisms from prokaryotes [42] to mammalian cells [24]. Trinucleotide repeat expansion diseases are therefore being evaluated for a potential contribution of nucleic acid structures to disease pathogenesis, with accumulating evidence that R-loops are involved in Fragile X syndrome [24, 43, 44] and Friedreich’s ataxia [44], with similar events also occurring in hexanucleotide repeat expansions [45]. We refer the reader to a number of excellent recent reviews of this topic for more complete insights into these unusual nucleic acid structures and their disease associations [13, 46, 47].

To establish a foundation for understanding their function, we mapped RNA:DNA hybrids genome-wide *in vivo* in two human cell lines with parallel transcriptional and proteomic studies. These studies provide new insights into how specific loci are preferentially selected as sites of formation of these structures, and allow the inference of some of their likely functional properties. These non-canonical nucleic acid structures occur in ribosomal DNA and at tens of thousands of loci in the remainder of the genome, with sequence characteristics indicating a polypurine-richness of the RNA in the hybrid that is likely to increase the thermodynamic stability of these structures. RNA:DNA hybrids appear to have heterogeneous and context-dependent properties, with subgroups showing relationships with local transcription and chromatin structural features, and a general trend towards decreased DNA methylation. On a more regional scale of hundreds of kilobases, RNA:DNA hybrids are enriched in regions of the genome with a greater abundance of L1 LINEs and CpG islands, and the chromatin modifications indicative of heterochromatin organization. These findings also support the possibility that the RNA generating these RNA:DNA hybrids is frequently generated *in trans,* a set of results that combines to provide new insights into these non-canonical nucleic acid structures in human cells.

## RESULTS

### RNA:DNA immunoprecipitation (RDIP)

We optimized an assay previously published as DNA:RNA immunoprecipitation (DRIP) [18] to map RNA:DNA hybrids, changing several components of the protocol. These updates include the pre-treatment of the cellular nucleic acid with RNase I, the use of sonication with the goal of minimizing bias in fragmenting the nucleic acid, and the addition of directional information about the strand derived from the RNA component of the hybrid. Given the extensive changes made, we distinguish the updated assay with the new acronym RDIP (RNA:DNA immunoprecipitation). The assay is based on the use of the S9.6 antibody, which is believed to recognize the intermediate A/B helical RNA:DNA duplex conformation, with little to no sequence specificity [48]. We performed extensive *in vitro* testing of the antibody to reconfirm these properties, including electrophoretic mobility shift assays and South-Western blots of oligonucleotides (including RNase H pre-treatment) that confirmed the necessary RNA:DNA hybrid specificity of the antibody (**Supplemental Figure S1a-e**).

The *in vivo* studies were focused on the primary, non-transformed, diploid IMR-90 lung fibroblast cell line because of the substantial genome-wide data available from the Roadmap Epigenomics Program [49]. For comparison, we isolated a clone of HEK 293T cells that we found to have the least copy number variability of several tested as determined by array comparative genomic hybridization (**Supplemental Figure S1f**). The immunoprecipitation using sonicated whole cell nucleic acid, pre-treated with RNase I, was optimized, and tested using a Southern dot blot using a (TTAGGG)_n_ probe to confirm enrichment of the telomeric TERRA-associated R-loop [34] (**Supplemental Figure S1g**). This pre-treatment with RNase I was recently shown to be necessary to reduce noise due to the S9.6 antibody detecting RNA in unusual conformations [2]. To allow the immunoprecipitated RNA:DNA hybrid to be ligated into sequencing adapters, an approach derived from RNA-seq library preparation was used. This provided the opportunity to introduce dUTP during second strand synthesis to reveal directional information about the strand on which the RNA molecule was located [50]. To confirm the RDIP-seq assay worked, we used peak calling analytical methodologies borrowed from ChIP-seq to identify the locations of RNA:DNA hybrids, followed by the use of single locus quantitative PCR to confirm enrichment in the immunoprecipitated material at these loci (**Supplemental Figure S1h**). Peaks were also verified at further loci using the orthogonal approach of bisulphite sequencing of non-denatured DNA to demonstrate the presence of the ssDNA that occurs at R-loops [51] (**Supplemental Figure S1i**).

### Subcellular localization studies

The subcellular localization of RNA:DNA hybrids has been studied in multiple organisms using a number of techniques [18, 26, 39, 52] and was investigated in the current study using two separate approaches. The first used limited amplification of the HEK 293T RDIP-seq library with a PCR primer to which the Texas Red fluorophore had been conjugated. This was hybridized to control human metaphases for visualization. As early results suggested that the pericentromeric region of chromosome 9 was generating signal, a locus-specific probe targeting the subtelomeric region of the p arm of this specific chromosome was included in the fluorescence *in situ* hybridization (FISH) study. **Figure 1a** depicts the results of these studies. A strong signal at the centromere of chromosome 9 is observed, as well as from the p arms of the acrocentric chromosomes, indicating enrichment at the Nucleolar Organising Regions (NORs), where ribosomal DNA (rDNA) repetitive sequences are located.

**Figure 1:**
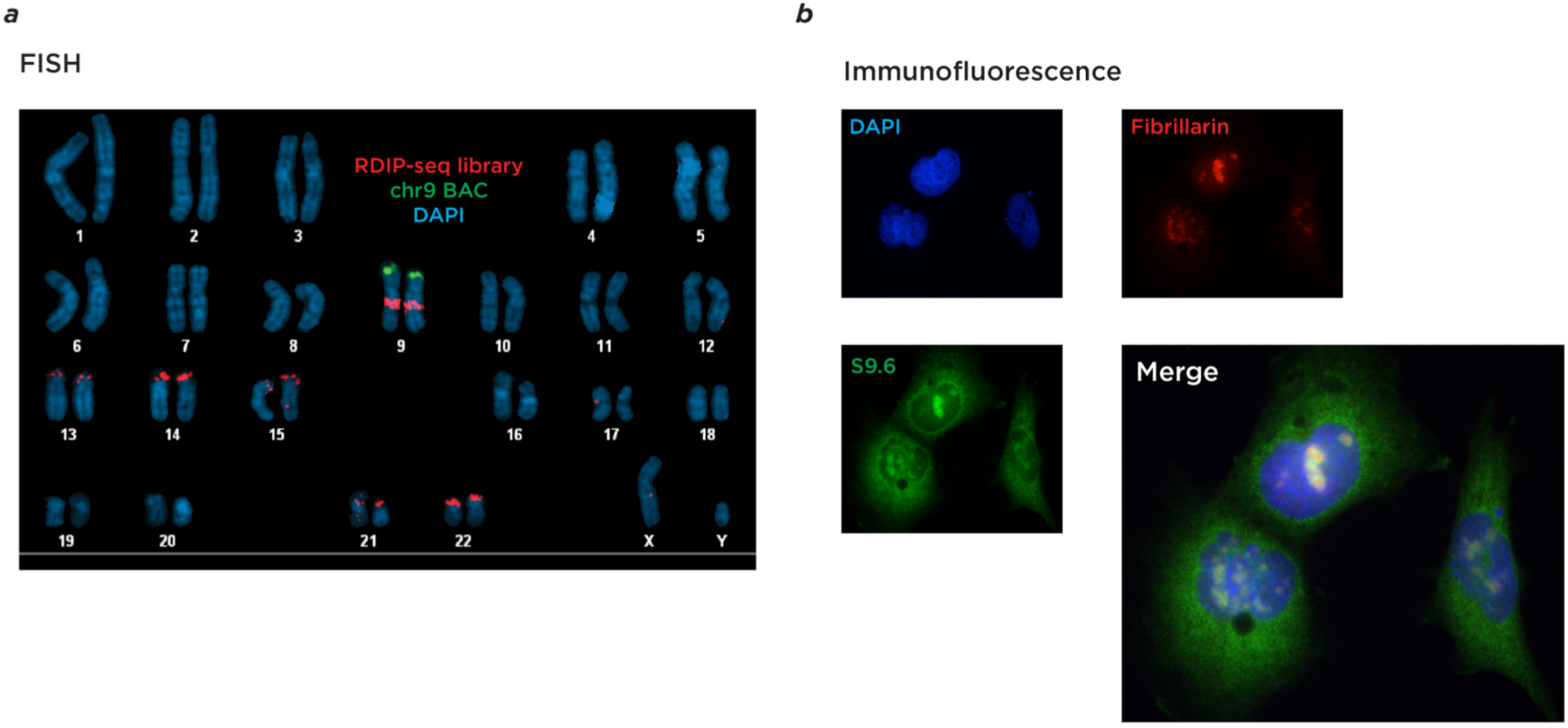
Subcellular localization studies. In panel (a) we show the results of hybridization of the fluorescently-labeled RDIP-seq library to a control male metaphase preparation. The RDIP-seq library is shown in red, a bacterial artificial chromosome (BAC) probe mapping to chromosome 9 in green, and DNA counterstained by DAPI in blue. We observe a specific strong signal from the RDIP-seq library mapping to the p arms of acrocentric chromosomes (HSA13-15 and HSA21-22), indicating enrichment at the nucleolar organizing regions (NORs) encoding ribosomal RNAs, and at the pericentromeric region of chromosome 9. In panel (b) we show the results of immunofluorescence using the S9.6 antibody (green) with an antibody to fibrillarin (red), demonstrating co-localization with the intranuclear S9.6 antibody signal (merge) and therefore enrichment in nucleoli. Further signal from the nuclear periphery and the cytoplasm using S9.6 is also observed, which may represent detection by this antibody of RNA conformations rather than RNA:DNA hybrids specifically [2].

The second subcellular localization approach employed was to use the S9.6 antibody for immunofluorescence of the HEK 293T cells. Consistent with previously published studies [18, 53], a subnuclear enrichment within nucleoli (confirmed with an anti-fibrillarin antibody, **Figure 1b**) was observed. Of note was the additional cytoplasmic signal that has also been noted in prior studies [53]. This signal may in part reflect signals from mitochondrial DNA [54] or the S9.6 antibody detecting ssRNA in unusual conformations [2].

### Ribosomal DNA studies

Prompted by the co-localization with the NORs seen in the subcellular localization studies, further investigation into RNA:DNA hybrid formation within ribosomal DNA was undertaken. The IMR-90 RDIP library was sequenced and mapped to a human reference genome including the consensus ribosomal DNA repeat unit [55] (accession number gi|555853|gb|U 13369.1|HSU13369), following the same approach as Zentner and colleagues [1]. The results showed that ~2% of reads mapped to the ribosomal DNA repeat unit and the remainder to the sequenced majority of the human genome. The mapping of reads to the rDNA repeat unit is shown in **Figure 2**. The immunoprecipitated RNA:DNA hybrids map heterogeneously within this repeat unit, with accumulation of reads at the known exons of the rDNA gene, and others in the intergenic spacer (IGS) region.

**Figure 2:**
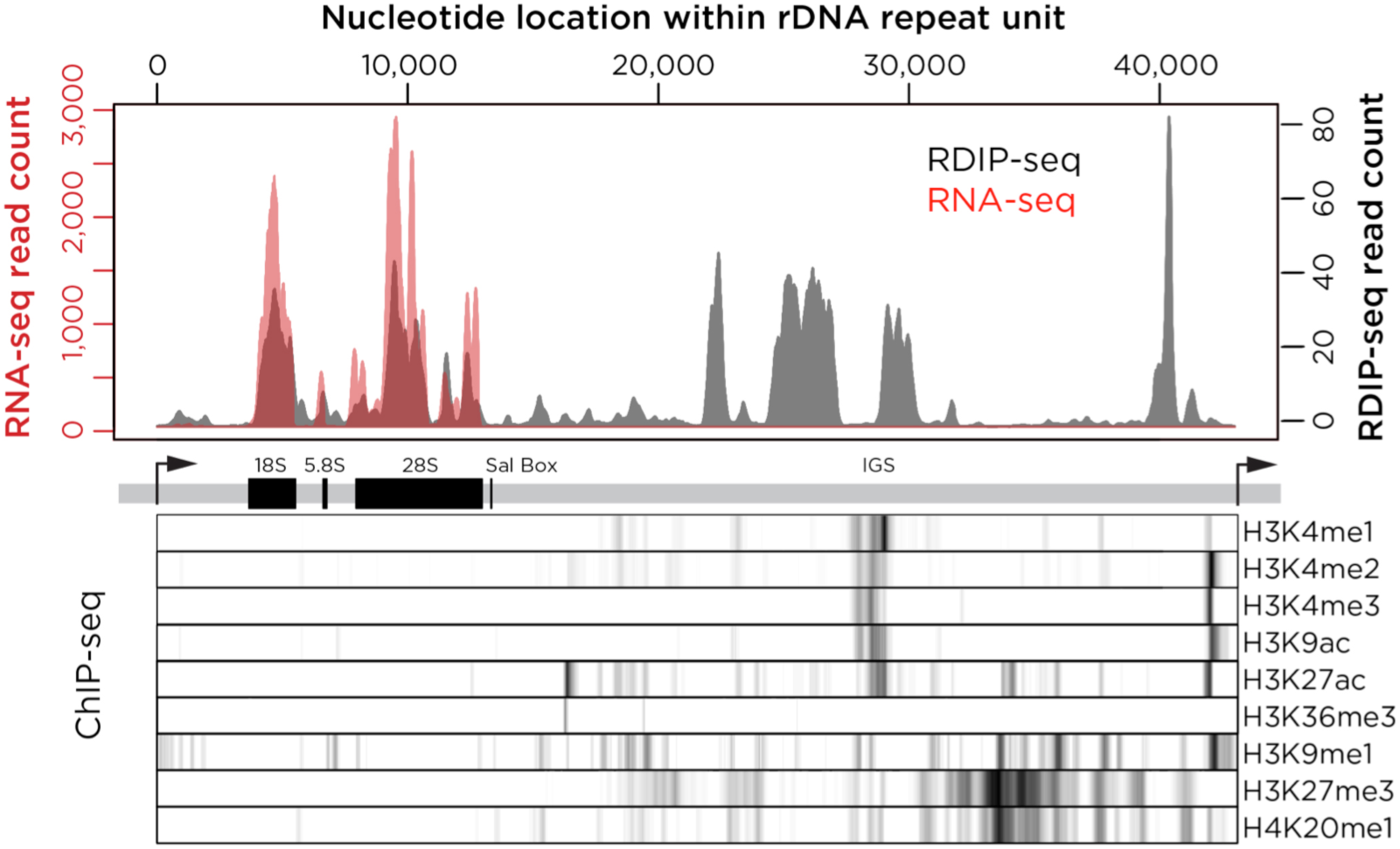
Mapping of RNA:DNA hybrids within the ribosomal DNA repeat unit. The upper panel shows the results of RDIP-seq (gray) and RNA-seq (red), with genomic annotations and results of ChIP-seq analysis in K562 cells [1] plotted below. RDIP-seq and RNA-seq data are both represented using a smoothed plot showing the number of reads aligned to each basepair of the repeating unit, while the ChIP-seq data signal intensity represents the mean value of non-overlapping 50 bp windows. RDIP-seq values were normalized by subtracting the frequencies of aligned reads of the input sample in each window. We find that RNA:DNA hybrids co-localize with the rRNA transcripts, but that there are also RDIP-seq peaks of comparable magnitude in the intergenic spacer (IGS) where no transcriptional activity is apparent from RNA-seq. The RNA:DNA hybrids in the IGS are upstream of the promoter region and flank the upstream candidate *cis*-regulatory sequence where there is H3K4 methylation and acetylation of H3K9 and H3K27.

To determine the relationship between the RNA:DNA hybrids and the transcribed sequences, RNA-seq on total RNA from the IMR-90 cells was performed without polyA selection or depletion of ribosomal RNA. This allowed deep sequencing of the expressed rRNA and colocalization with the RNA:DNA hybrid reads (**Figure 2**). The RDIP-seq reads in the 5’ end of the repeat unit are precisely co-localized with the RNA-seq reads, but there is RNA:DNA hybrid formation with comparable read enrichment in the IGS region. Using K562 cell ChIP-seq data provided by Zentner and colleagues [1], the RNA:DNA hybrids are found to be located upstream from the rDNA promoter and flanking the candidate *cis*-regulatory sequence in the IGS region (**Figure 2**). The intergenic candidate *cis*-regulatory sequence was also shown to occur in embryonic stem cells, umbilical vein cells and normal human epidermal keratinocytes [1], and thus appears to be constitutive. It is therefore reasonable to predict that the element is also present in the IMR-90 cells. Some of the rDNA RDIP-seq signal is attributable to RNA:DNA hybrid formation involving the canonical rRNA transcript, but further RNA:DNA hybrids are formed in the IGS ribosomal DNA region sparing the regions containing candidate *cis*-regulatory elements.

### Genome-wide studies

Having defined the source of the rDNA signal, the focus turned to the majority of reads that mapped to the remainder of the sequenced genome. There are tens of thousands of RNA:DNA hybrid-forming loci (mapped as peaks using a ChIP-seq analytical approach) throughout the human genome (**Figure S2**), the same magnitude observed previously in DRIP-seq experiments [18]. There is a significant enrichment for loci shared by IMR-90 and HEK 293T cells, indicating that many RNA:DNA hybrid-forming loci may be constitutive across cell types. Focusing on the loci in the human diploid IMR-90 fibroblast cell line, RNA:DNA hybrids are demonstrated to be distributed genome-wide, with most of the peaks located in intergenic regions (**Figure 3a**). The enrichment of peaks in each of these major genomic contexts was calculated and the significance of enrichment was tested based on overlap (nucleotide occupancy) using permutation analyses. **Figure 3b** shows that promoters (and the highly correlated CpG island feature) are strongly enriched for RNA:DNA hybrids, and that they are distributed elsewhere in the genome at close to expected frequencies, apart from a modest but significant depletion at RefSeq gene bodies and intergenic regions (excluding promoter and lncRNA sequences).

**Figure 3:**
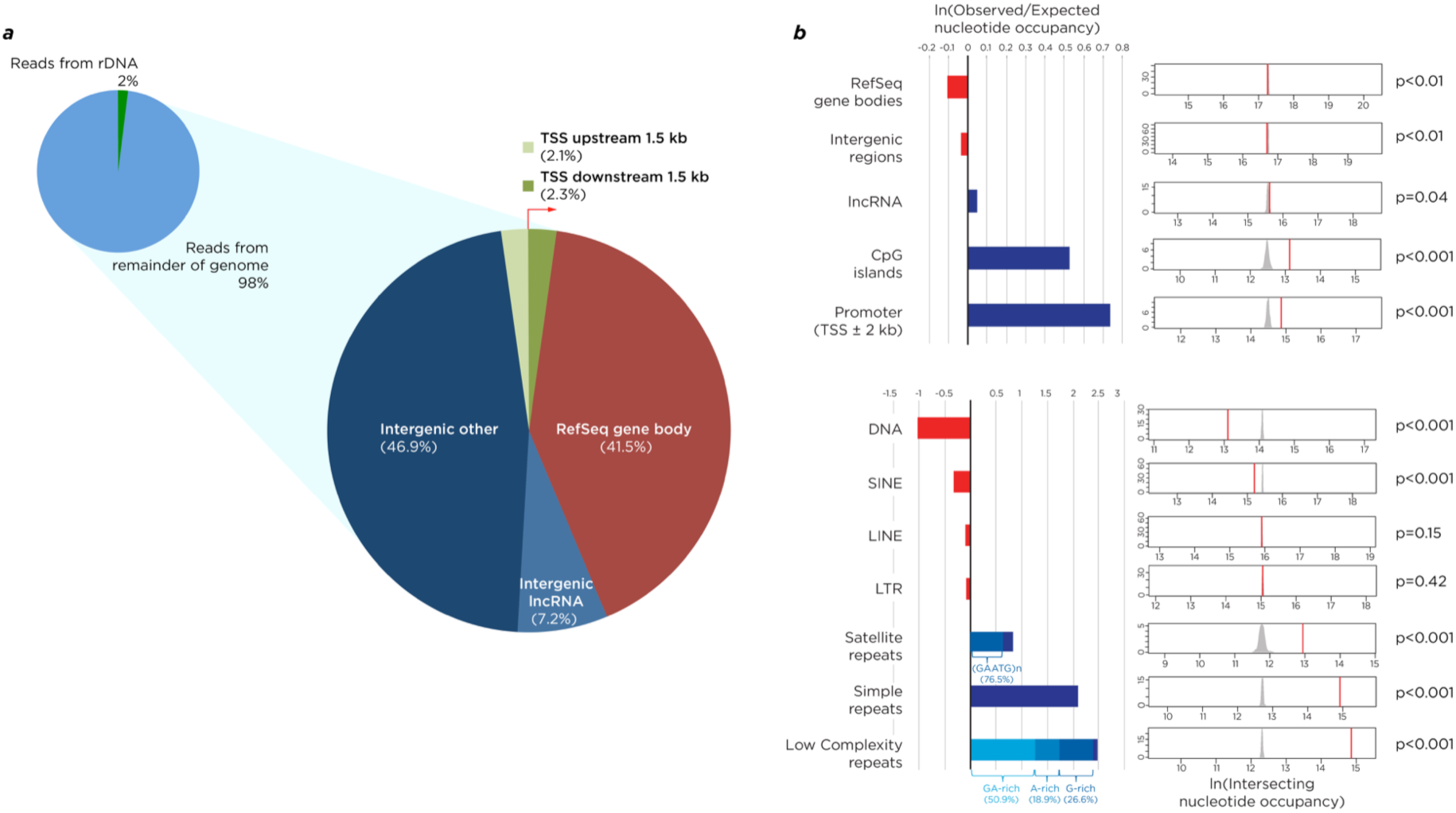
Genomic distribution of RNA:DNA hybrids. In panel (a) we show that the proportion of reads mapping to rDNA is 2%, and break down the remaining 98% by genomic context, showing the majority of RNA:DNA hybrids (called as peaks using ChIP-seq analytical approaches) to be located in intergenic regions. To understand these RNA:DNA hybrid distributions, we calculated observed/expected ratios based on nucleotide occupancy of genomic features, and performed permutation analyses testing for the likelihood of randomized intersection (b), the results of which are shown in **Supplemental Table S1**. We found depletion of RNA:DNA hybrids at RefSeq gene bodies, intergenic regions, and SINE and DNA transposable elements but significant enrichment at promoters and CpG islands, and a number of purine-rich repetitive sequences.

As RNA:DNA hybrids in yeast have been shown to be enriched at transposons [30], their representation within sequences annotated as repetitive within the human genome was explored. In **Figure 3b**, the sequences annotated as low complexity and simple repeats by RepeatMasker are shown to be the most strongly over-represented, but satellite repeats are also found to be enriched in RNA:DNA hybrids. When the low complexity repeats were explored in greater detail, the strand on which the RNA component of the RNA: DNA hybrid was located was found to be composed of GA-rich, G-rich, and A-rich families of low complexity repeats. Additionally, within the satellite repeats that co-localized with the RNA of RNA:DNA hybrids, 76.5% of the repeats were (GAATG)_n_ sequences.

It is known that purine-rich RNA binds *in vitro* with greater affinity to its pyrimidine-rich DNA complement than the equivalent purine-rich DNA sequence [14, 22], which may indicate a role for biochemical stability maintaining RNA:DNA hybrids *in vivo.* As the analyses of repetitive sequences suggested enrichment of purine-rich RNA in these RNA:DNA hybrids, this finding was explored more fully, testing for and finding from the genome-wide data a strong intramolecular skewing towards GA:CT enrichment (**Figure 4a**). To test globally whether this purine (GA) enrichment was present on the RNA-containing strand, the directional sequence information was used to examine nucleotide skewing on each strand at RNA:DNA hybrids, confirming the RNA-derived sequence to be strongly purine-enriched (**Figure 4b**). The 10% of peaks with the least tendency towards having the RNA enriched on one strand were removed from further analyses as being likely to over-represent experimental noise.

**Figure 4:**
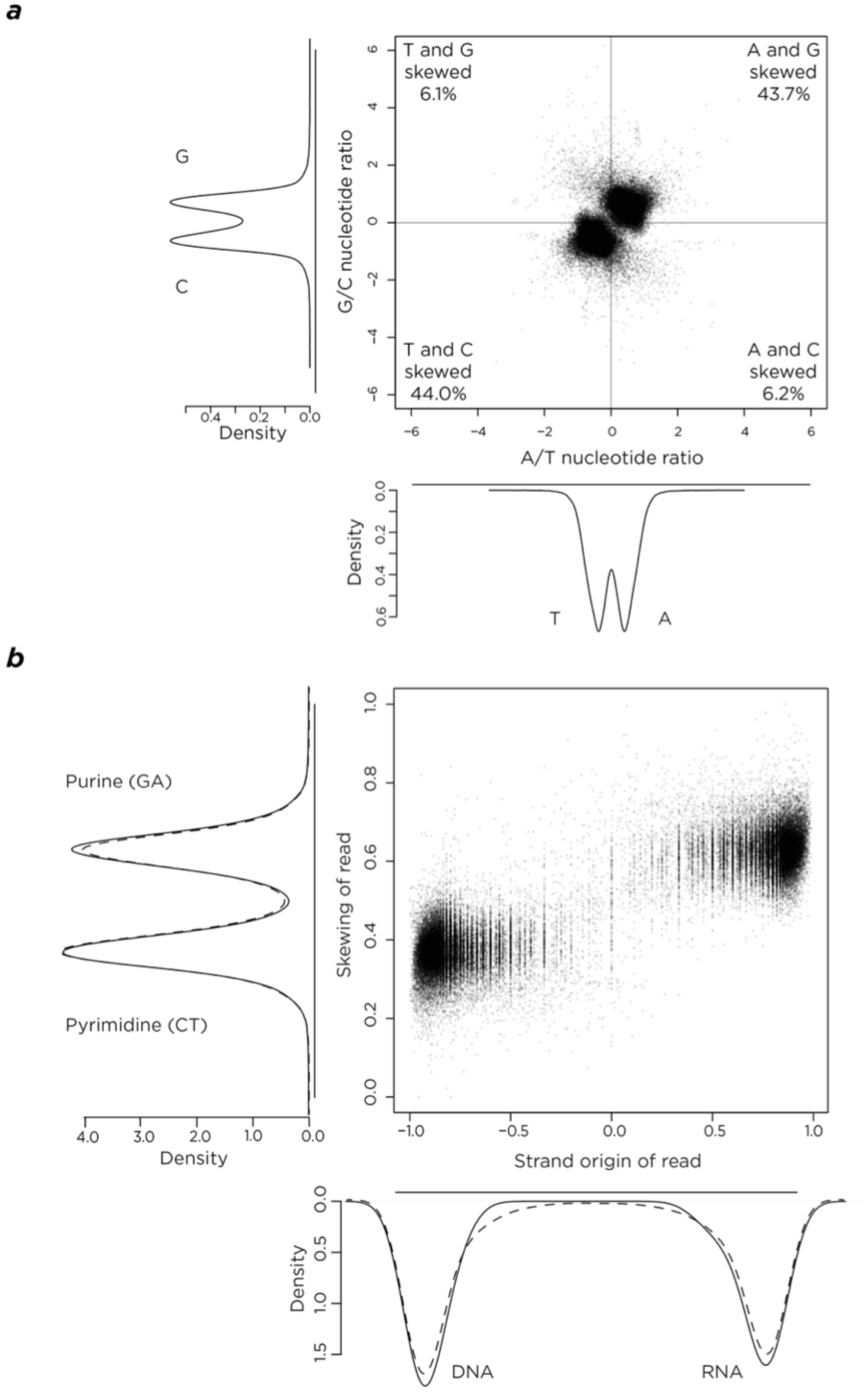
Nucleotide skewing analyses. In panel (a) we plot the skewing within a strand of A compared to T (x axis) or G compared to C (y axis) in the RNA:DNA hybrid peaks genome-wide. We find that the peaks are strongly over-represented for purine (G+A) and pyrimidine (C+T) skewing. As our sequencing approach allowed us to identify the RNA and DNA-derived strands separately in the RNA:DNA hybrid, in (b) we proceeded to test whether there was a relationship between skewing (based on the number of G+A divided by the total number of nucleotides) and each type of nucleic acid-derived sequence, finding a clear enrichment for purine skewing on the RNA-derived strand.

### Relationship of RNA:DNA hybrids to local transcription

As some RNA:DNA hybrids have been found to have transcriptional termination properties [56, 57], it was tested whether the RDIP directional sequencing allowed the observation of the an orientation bias within genes. This tendency has been observed for transposable elements, which are believed to have different effects on gene function depending on their insertion orientation in gene bodies [58, 59]. The nucleotide skewing within each peak was visualized, revealing the purine-enriched component to be displaced 5’ from the mid-point of the peak (**Figure S3**), which is consistent with the RDIP protocol using the RNA component of the RNA:DNA hybrid to prime second strand synthesis, proceeding unidirectionally 3’ and relatively under-representing the region 5’ to the RNA. This observation is independently supportive of the RNA component of the RNA:DNA hybrid being purine-enriched. There was a modest orientation bias against purine-rich sequences in the same orientation as the gene (**Figure S3b**), indicating that most but not all genes tolerate an RNA:DNA hybrid with the RNA on the transcribed strand.

To explore the relationship between RNA:DNA hybrid formation and transcription further, the proportions of genes with peaks were tested for transcription states from the RNA-seq data, finding that most transcribed RefSeq genes do not contain RNA:DNA hybrids but that the transcribed genes have a higher frequency of RNA:DNA hybrids than non-transcribed genes (7.75% compared with 6.09%, **Figure 5a**). The locations of these RNA:DNA hybrids within genes was defined using a metaplot, identifying the first ~1.5 kb downstream from the transcription start site (TSS) as the region most consistently enriched (**Figure 5b**). This region is also found to be modestly enriched in purine skewing for genes with and without RNA:DNA hybrids (**Figure 5c**). Surprisingly, given the transcriptional termination properties attributed to RNA:DNA hybrids [56, 57], the transcriptional end site is notable for a slight depletion of these structures (**Figure 5b**). The information from lncRNAs also suggests a modest enrichment for RNA:DNA hybrids in the immediate vicinity of the TSS (**Figure S4**). The local generation of RNA:DNA hybrids has previously been described to be associated with transcription of the region [16, 24], so the genes were stratified by expression level, finding that the proximal 1.5 kb region downstream from the TSS showed an increase in peaks associated with increasing quantiles of gene expression states (**Figure 5d**). The conclusion is that transcriptional levels have effects on the likelihood of forming RNA:DNA hybrids, and that local purine enrichment may increase the tendency of these structures to be formed in the ~1.5 kb immediately downstream of the TSS in a small subset of genes.

**Figure 5:**
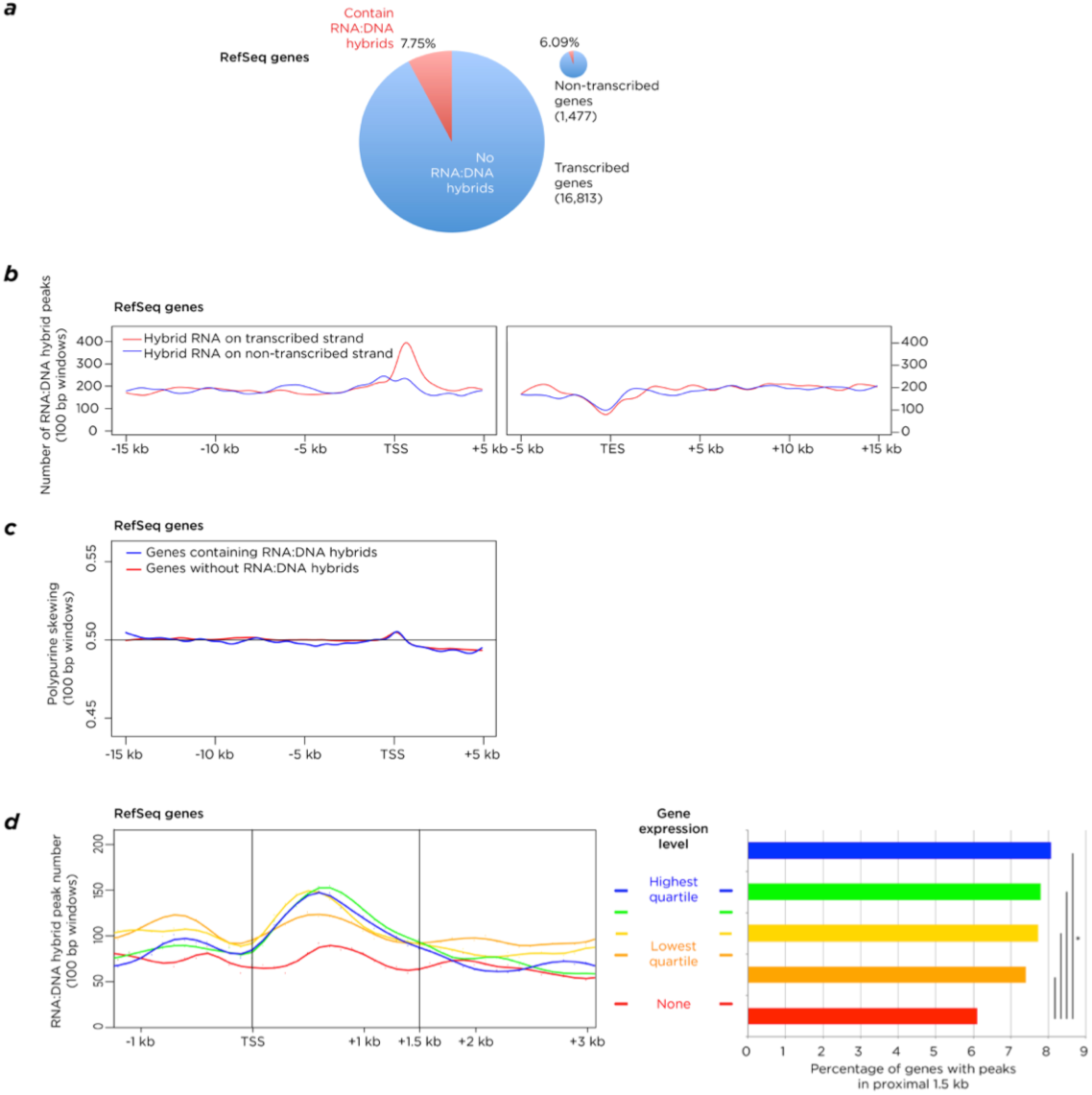
Transcriptional relationships of RNA:DNA hybrids. In (a) the proportion of RNA:DNA hybrid peaks in transcribed genes is shown to be higher than in nontranscribed genes, but that the majority of genes do not contain RNA:DNA hybrids. In (b) a metaplot of RNA:DNA hybrid peaks is shown, illustrating the number of peaks intersecting with 100 bp windows, with the RNA of the hybrid on the transcribed strand of the gene (red) or the opposite strand (blue). This revealed an enrichment of the RNA-derived sequence on the transcribed strand in the first ~1.5 kb downstream from the transcription start site (TSS). A depletion of RNA:DNA hybrids is found at the transcription end site (TES). In (c) we show that the region immediately downstream from the TSS is purine-skewed, represented by skewing values of 100 bp windows averaged for all genes, but that this is to the same degree in genes that form RNA:DNA hybrids (blue) as those genes that do not form these structures (red). In (d) a metaplot of RefSeq genes (left) shows that the transcription level of genes (as measured by RNA-seq) is positively associated with the number of RN:DNA hybrids intersecting with 100 bp windows immediately downstream of the TSS. This reflects only modest increases in the small proportions of genes forming peaks (right), though found to be a significant relationship using a proportions test.

As RNA-seq measures the steady state of RNA in the cell, and does not necessarily reflect active transcription, we added an analysis of global run-on sequencing (GRO-seq) data generated previously for IMR-90 cells [60]. We found 24,875 RNA:DNA hybrids in these cells to map to RefSeq genes, 24,174 to active genes of which 23,717 had evidence for transcription from GRO-seq data. Interestingly, of the 788 RNA:DNA hybrids mapping to genes with no evidence for transcriptional activity from our RNA-seq data, 733 mapped to loci where GRO-seq indicated local transcription. Exploration of the 57,308 RNA:DNA hybrids mapping to intergenic regions revealed 11,825 to map to loci where GRO-seq indicated transcription. These data suggested GRO-seq to be a more sensitive indicator of transcription than RNA-seq or genomic context, prompting us to explore how many of the RNA:DNA hybrids mapped to loci where GRO-seq indicated transcription, finding evidence for transcription at 27,352 (47.7%) of the RNA:DNA hybrids.

### Relationship of RNA:DNA hybrids to regulators of transcription

To begin to infer any transcriptional regulatory function of the RNA:DNA hybrids from their genomic locations, studies were performed correlating RNA:DNA hybrid locations with enrichment or depletion for other chromatin and transcriptional regulators directly overlapping the RNA:DNA hybrids. Using IMR-90 bisulphite sequencing data from the Roadmap Epigenomics Project (accession number NA000020923.1), a modest decrease in DNA methylation within RNA:DNA hybrids was found compared with genome-wide levels, a finding which is consistent with the hypomethylation of DNA previously observed for RNA:DNA hybrids at CpG islands [18] (**Figure S5a**). *In vitro* studies have shown RNA:DNA hybrids to be refractory to the formation of nucleosomal structures [19], a finding supported by the observation that 7.46% of all RNA:DNA hybrids overlap DNase hypersensitive sites, representing a significant association genome-wide (**Figure S5b-c**). We also tested whether there may be unrecognized transcription at enhancer RNAs [61, 62] at sites of formation of RNA:DNA hybrids. We found that of the 57,308 RNA:DNA hybrids, 4,277 (7.5%) map to DNase hypersensitive sites, of which 3,560 have the H3K27ac modification, most of which (3,434) also have the H3K4me1 modification. Of these H3K27ac/H3K4me1 DNase hypersensitive sites with RNA:DNA hybrids, 2,154 have evidence for transcription from GRO-seq. As enhancer RNAs are inherently unstable [63], it is possible that transcription is actually occurring at a higher proportion of these loci that the GRO-seq data would indicate, potentially linking transcription with RNA:DNA hybrid formation at as many as the 7.5% located at DNase hypersensitive sites. A motif analysis of RNA:DNA hybrid-forming loci genome-wide revealed an enriched polypurine (GGAA)_n_ sequence, which has been associated with binding by the FLI1 transcription factor [64] (**Figure S6a**).

A notable macro-scale organization of RNA:DNA hybrids was apparent in the human genome, with regions of dense and sparse RNA:DNA hybrid formation (example shown in **Figure S6b**). Using publicly-available ChIP-seq data from the IMR-90 cell line, it was possible to ask whether RNA:DNA hybrids in the human genome occur in regions of distinctive regulatory characteristics. We have previously noted that there is extensive inter-correlation of genomic features [65], making it difficult to discriminate specific associations when there are multiple correlating genomic variables. In order to explore the transcriptional and regulatory context of RNA:DNA hybrid peaks, regression models were fitted to the data, regularized using the least absolute shrinkage and selection operator (LASSO; [66]) with the peak density as the response variable. Least angle regression (LARS; [67]) was used, progressively adding covariates to the model and testing the significance of each added predictor using the covariance test statistic proposed by Lockhart *et al.* [68]. The results of this procedure are shown in **Figure 6**. The first covariate to enter the model as significantly enriched in co-localization with RNA:DNA hybrids in 500 kb windows is the repressive histone mark, H3 lysine 27 trimethylation (H3K27me3), followed by CpG islands, L1 LINE retroelements and a further repressive histone mark, H3K9me3. The first eight covariates to enter the model all gave significant values of the covariance test statistic.

**Figure 6:**
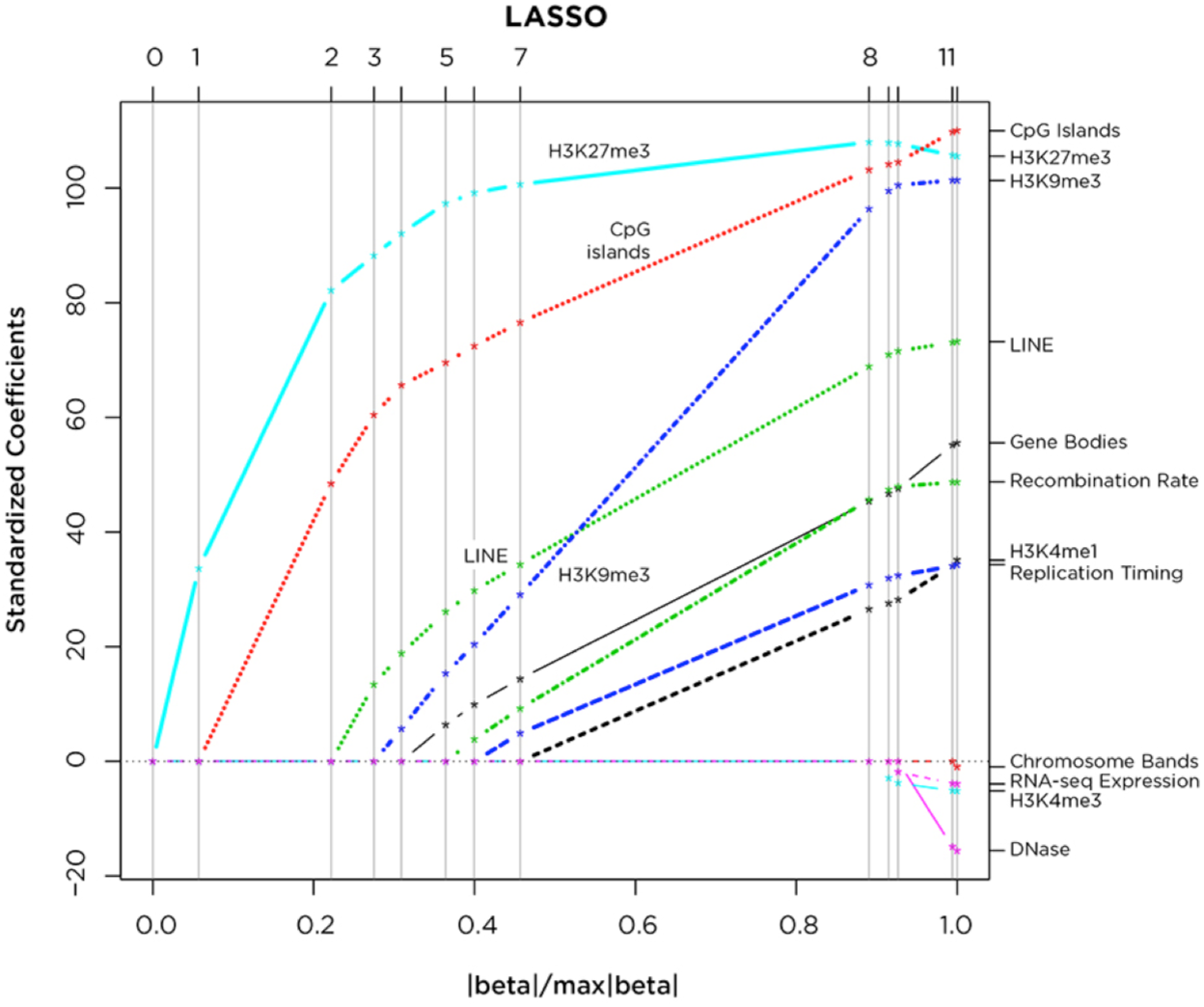
Macro-scale genomic associations of RNA:DNA hybrids. We used a least absolute shrinkage and selection operator (LASSO) adaptive regression approach to explore the association of genomic sequence features with RNA:DNA hybrid density in 500 kb windows. The figure shows the order in which covariates enter the model as the constraint on the sum of the regression coefficients (x-axis) is progressively relaxed from 0 to its maximum value (corresponding to the ordinary least squares regression vector).

### Local chromatin organizational studies using mass spectrometry

Finally, characterization of chromatin located at RNA:DNA hybrids was performed to identify the proteins enriched at these loci. Chromatin from HEK 293T cells was sonicated and a fraction immunoprecipitated with the S9.6 antibody, eluting the protein complexes using RNA:DNA hybrid oligonucleotides, and identifying local proteins through mass spectrometry (**Figure 7a**). These results and Western blotting validation of candidate proteins of interest are shown in **Figure 7b** and **Supplemental Table S2. A** number of different specific proteins plausibly associated with RNA:DNA hybrids were identified. RNA helicase A (encoded by *DHX9*) is a protein known to be involved in resolving RNA:DNA hybrids [69] and is a necessary partner for FLI1 in tumourigenesis [70], while DNA binding protein B (YBX1) is known to bind to ssDNA [71] which should be part of R-loops formed at these loci. ILF2 and ILF3 are also found in the chromatin at RNA:DNA hybrids. These are transcription factors known to recognize a purine-rich motif [72], with our results raising the possibility that their binding may depend on the target nucleic acid existing in an RNA:DNA conformation.

**Figure 7:**
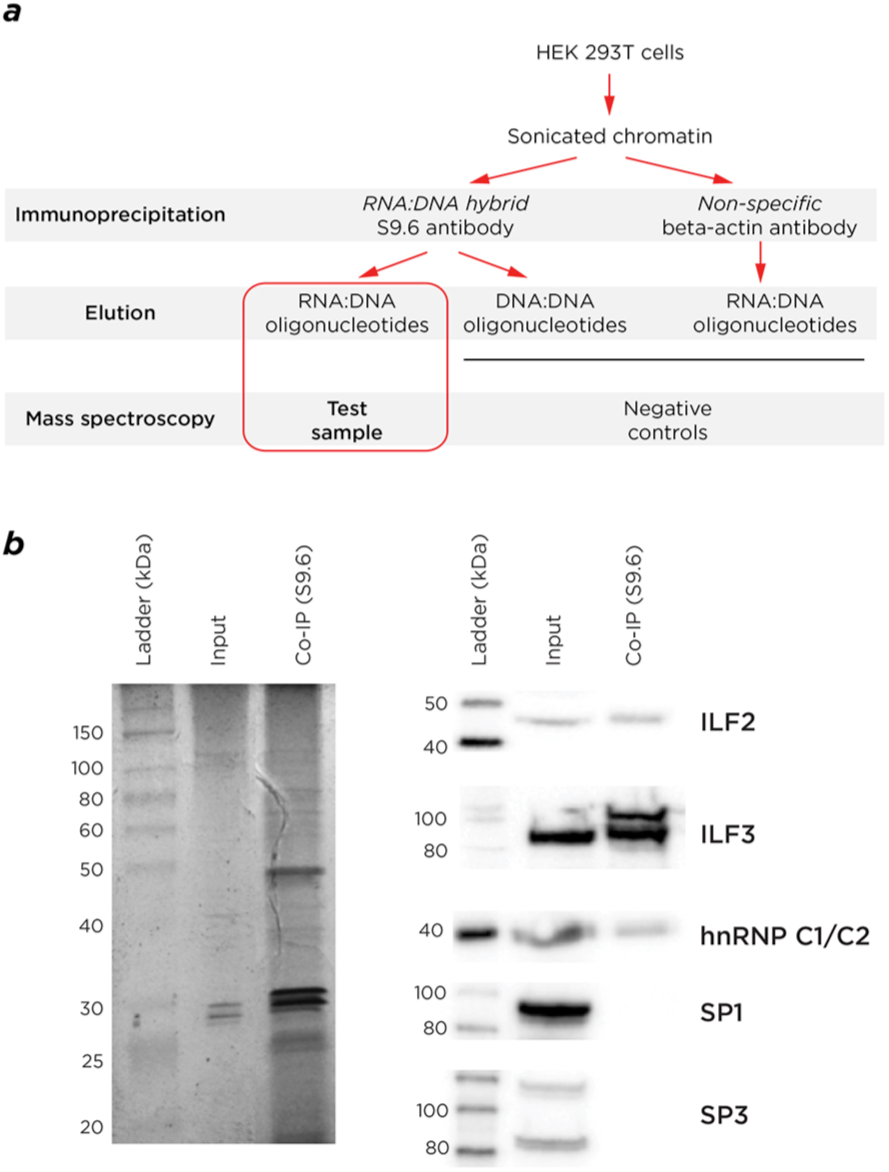
Chromatin organizational studies at RNA:DNA hybrids using mass spectrometry. In panel (a) we show the experimental approach used for these proteomic studies. In (b) the altered pattern of enriched proteins compared with the input sample is seen using gel electrophoresis, and the results of Western blots confirming the enrichment of specific candidate proteins identified by mass spectrometry (ILF2, ILF3, hnRNP C1/C2), with SP1 and SP3 as controls known to bind to G-skewed DNA motifs.

The presence in local chromatin of RNA helicases and topoisomerases is consistent with prior reports that these enzymes are involved in the removal of RNA:DNA hybrids [28, 44]. The question arose whether the IMR-90 and HEK 293T cells express the genes encoding the broader group of proteins implicated in removal of RNA:DNA hybrids *in vivo.* Using the RNA-seq data, nine of these genes were categorized into quartiles of expression, finding that all of the genes were expressed at high levels (**Figure S7**). The presence of the RNA:DNA hybrids in these cells is therefore in spite of robust levels of expression of genes encoding proteins that should actively remove them.

## DISCUSSION

Mapping RNA:DNA hybrids in human cells has allowed new insights into the properties of these non-canonical nucleic acid structures. We confirm through subcellular localization studies prior observations that the ribosomal DNA harbors these structures [18] (**Figure 1**). Additionally, we expand on findings in yeast [30] by mapping RNA:DNA hybrid locations within the human rDNA repeating unit, revealing these structures to be formed not only at the expressed rDNA gene but also in the intergenic spacer sequence (**Figure 2**). The signal from this repetitive sequence is necessarily composed of all rDNA repeat units in the genome, so we cannot distinguish events occurring within individual alleles, but we can make several inferences. Firstly, that the enrichment of RNA:DNA hybrids within the rDNA repeat unit is not uniform but is enriched at two types of loci, the exons of the rRNA genes and the intergenic spacer sequence where they spare present candidate *cis*-regulatory loci (**Figure 2**). The mapping of RNA:DNA hybrids to the rDNA gene exons is an interesting finding as it implies that the RNA associating with the rDNA is already spliced and not the primary transcript through the region. This is less supportive of a co-transcriptional model for RNA:DNA hybrid formation [31, 32] and more indicative of rRNA acting *in trans* to generate these structures, as has been found for RNA:DNA hybrids in yeast [26].

The mapping of reads to the rDNA repeat was consistent with the imaging data indicating the presence of RNA:DNA hybrids in nucleoli (**Figure 1**), allowing us to proceed with confidence to assess the distribution of the majority of the reads elsewhere in the genome. The first observation was that the RNA:DNA hybrids were not enriched in gene bodies relative to intergenic sequences (**Figure 3a-b**), again failing to support their presence being solely a function of recognized transcription. Furthermore, the rRNA model would suggest that spliced mRNAs might associate *in trans* with their genes of origin, but this is not reflected by overrepresentation of RNA:DNA hybrids in RefSeq genes (**Figure 3b**). Instead we observe that a small proportion of genes have peaks within their bodies (**Figure 5a**), with a significantly higher proportion of expressed genes than silent genes containing RNA:DNA hybrids (**Figure 5d**). These tend to form in the ~1.5 kb immediately downstream of the transcription start site, where they are influenced by the level of transcription (**Figure 5d**) but can be found even in genes that are not measurably expressed by RNA-seq (**Figure 5a,d**), and are overall depleted in RefSeq gene bodies (**Figure 3b**). Analysis of GRO-seq data from IMR-90 cells [60] reveals evidence for transcriptional activity at genes categorized as silent by RNA-seq, and at loci of open chromatin in the genome, which should include sites of transcription of enhancer RNAs [61, 62]. Overall, however, even these sensitive indicators of active transcription only account for a subset of RNA:DNA hybrids in the genome, indicating that transcription through a locus is therefore only moderately influential in generating these structures.

Adding to the tendency of the proximal 1.5 kb to form RNA:DNA hybrids is the enrichment at this location genome-wide for purine-skewed DNA in the transcriptional orientation of the gene (**Figure 5c**). We first noticed that purine enrichment may be a property of RNA:DNA hybrids *in vivo* when we found a strong enrichment for repetitive sequences composed of polypurines in our RepeatMasker analysis (**Figure 3b**). We confirm the purine skewing to be a general property of these sequences (**Figure 4** and **Figure S3**), which extends prior observations that suggested isolated G density [16] or GC [21] skewing, to be characteristic of these loci. As purine-rich RNA binds to complementary pyrimidine-rich DNA with greater affinity than the same purine-rich DNA sequence *in vitro* [14, 22], this is likely to be a factor in the ability of the RNA to maintain displacement of the ssDNA in the R-loop structure.

While transcriptional termination has been described to be a property of RNA:DNA hybrids [56] (reviewed in [13]), we observe that RNA:DNA hybrids are not enriched at the annotated ends of RefSeq genes and are, in fact, relatively depleted (**Figure 5b**). However, we see a small orientation bias in RefSeq genes, with a shift away from RNA:DNA hybrids with the RNA in the same orientation as transcription (**Figure S3**). We interpret this to indicate that a subset of RNA:DNA hybrids may cause transcriptional disruption effects, but that it is not a universal property throughout the genome.

We can infer some likely functional properties of RNA:DNA hybrids by genomic co-localization and proteomic approaches. The genomic co-localization studies were both immediately at the RNA:DNA hybrid location and more broadly in their flanking regions, the latter prompted by what appeared to be higher-scale organization of the distribution of these loci (**Figure S6b**) and by prior studies in yeast [36]. The immediate local features included DNase hypersensitivity (**Figure S5b-c**), which is consistent with prior *in vitro* published findings that nucleosomes do not readily form on these structures [19]. The tendency of RNA:DNA hybrids to be resistant to acquisition of DNA methylation [18] finds some support from our data, but the modest degree of relative hypomethylation indicates that the effects occur at only a small subset of loci. In the regional analysis of the co-localization of RNA:DNA hybrids and genomic sequence features within 500 kb windows of the genome, the enrichment found for CpG islands was not surprising given our observations that promoter-proximal sequences are enriched in RNA:DNA hybrids (**Figure 3b**). However, the enrichment in the same broader regions for the repressive H3K27me3 and H3K9me3 marks was unexpected for structures with the possibility of being co-transcriptionally generated. We interpret this to indicate one of the following three possibilities: that these regions are more transcribed than we can appreciate using the data available to us, allowing co-transcriptional formation of RNA:DNA hybrids, or that RNA forming RNA:DNA hybrids *in trans* is better able to target these regions, or that these structures are more stable in the context of repressive heterochromatin, with a causal model prompted by observations in fission yeast [36] that would involve the RNA:DNA hybrids having a mechanistic role to induce the regional repressive organization.

The proteins revealed by the proteomic studies were consistent with the local presence of RNA:DNA hybrids and R-loops (**Figure 7, Supplemental Table S2**), including RNA helicase (DHX9) and single-stranded DNA binding properties. We were especially intrigued by the presence of the ILF2 and ILF3 components of the Nuclear Factor of Activated T-cells (NF-AT) transcription factor, which is required for T-cell expression of interleukin 2 and represents a target of the immunosuppressive Cyclosporin A and FK506 drugs [73]. ILF2 (NF45) and ILF3 (NF90) are characterized by their binding to polypurine-rich interleukin gene enhancers [72], and are described to have the property of being able to bind to dsRNA *in vitro* [74]. This property, when combined with our finding of enrichment in chromatin at RNA:DNA hybrids, suggests that the selective binding of NF-AT at specific genomic locations may be dependent upon those sites being in an RNA:DNA hybrid conformation, which is structurally more similar to A-form dsRNA than B-form dsDNA [14]. The sequence motif (GGAA)_n_ that we found to be enriched at RNA:DNA hybrids (**Figure S6a**) closely resembles that of the FLI1 transcription factor [75]. FLI1 is a master regulator of hematopoiesis [76] in the ETS family, and has been causally implicated in pediatric Ewing’s sarcoma [77]. The oncogenic effect of FLI1 (as a fusion protein with EWS) is enhanced by RNA helicase A [70] which it appears to inhibit [78], an interaction that can in turn be inhibited by small molecules with therapeutic potential [79]. Expression of EWS-FLI1 induces chromatin opening at sequences with the (GGAA)_n_ motif [80]. The combination of the findings of binding to a polypurine-rich motif and interaction with RNA helicase A combine to suggest that FLI1 may also bind to an RNA:DNA nucleic acid conformation.

The model for RNA:DNA hybrid physiology that results from our studies indicates that they form as a result of an equilibrium between formation, stability and removal, with increased transcription having only a modest influence for the small subset we believe to be formed co-transcriptionally. Once formed, those at purine-skewed loci are likely to be more stable thermodynamically, while the presence of enzymes like RNA helicase A in the local chromatin and the robust expression of genes encoding proteins that remove RNA:DNA hybrids (**Figure S7**) reflect how these structures remain despite active processes dedicated to their removal. The RNA:DNA hybrids form DNase hypersensitive structures which may facilitate or reflect binding of transcription factors with preferences for either the A/B form RNA:DNA duplex or the ssDNA in the R loop, and exist in large scale domains of repressed chromatin, with which their causal relationship is uncertain. We propose that the weight of evidence supports many of the RNA:DNA hybrids being formed *in trans,* by RNA transcripts originating from regions of the genome other than the location of the RNA:DNA hybrid itself. The ability of RNA to invade a double stranded DNA molecule *in trans* is being strikingly highlighted at present by CRISPR/Cas technology, which creates an RNA:DNA hybrid as part of an R-loop [81]. We find little evidence for the majority of the RNA:DNA hybrids *in vivo* to be located at recognizably transcribed sequences. More persuasively supporting a *trans* hypothesis is the finding that the RNA:DNA hybrids in the rDNA repeat unit map to processed rather than primary rRNA transcripts. The simplicity of the polypurine-skewed sequences at RNA:DNA hybrids potentially allows a limited number of transcripts to target a large number of loci. The nuclear-retained polypurine-rich RNAs found in mammalian cells represent a type of non-coding RNA of unclear function [82] that could mediate such *trans* effects *in vivo.* Overall, it appears that there are numerous influences upon physiological RNA:DNA hybrid formation, the dissection of which will be essential if we are to understand the roles ascribed to them in disease states [83].

## CONCLUSIONS

A systematic analysis of RNA:DNA hybrids in human cells reveals their presence throughout the genome, including in the ribosomal DNA repeat unit, cumulatively representing millions of base pairs of DNA. The results help to resolve a number of conflicting theories about the formation of RNA:DNA hybrids, with only small influences of local transcription found, and evidence indicative of their formation *in trans.* Their sequence characteristics are clearly shown to be defined by purine enrichment for the RNA component of the hybrid, supporting a thermodynamic characteristic of RNA:DNA hybrids that should favor their stability. Functionally, we find evidence linking the presence of these structures to local DNA methylation and local and regional chromatin organizational states, with proteomic studies revealing the presence of transcription factors that may be binding preferentially to the RNA:DNA conformation. The contribution of non-canonical nucleic acid structures in transcriptional regulation is underexplored but warrants further investigation, adding a new layer of information in understanding transcriptional regulation in mammalian cells.

## METHODS

### S9.6 antibody production

The S9.6 antibody-producing hybridoma line was purchased from ATCC (HB08730), and the hybridoma line was grown in Integra Flasks by our institution’s monoclonal antibody core facility in serum-free medium. The S9.6 antibody was then purified by the macromolecular therapeutics core facility using a Protein-G column and size exclusion. The antibody was validated using an electrophoretic mobility shift assay (EMSA) and southwestern blotting to test for specificity to RNA:DNA hybrid oligonucleotides. A full description of these experiments is provided in the **Supplemental Experimental Procedures**.

### Immunofluorescence

HEK 293T cells were fixed in 4% paraformaldehyde for 10 minutes at room temperature, and then permeabilized for 10 min with 0.5% Triton-X-100. The cells were immunostained with anti-S9.6 antibody and anti-Fibrillarin antibody (Cell Signaling) for 1 hour, washed three times with phosphate buffered saline (PBS), and incubated with Alexa Fluor 488-labeled anti-mouse IgG antibody and Alexa Fluor 568 labeled anti-rabbit IgG antibody (Invitrogen) for 30 minutes at room temperature. Finally, cells were mounted in mounting solution ProLong Gold with DAPI (Invitrogen).

## FISH

Fluorescence *in situ* hybridization (FISH) was performed using our previously published approach [84]. For the experiment described, 2 *μ*g of DNA from the Illumina RDIP-seq library were labeled by nick translation using spectrum orange-dUTP (Invitrogen, Carlsbad, CA). A locus-specific BAC clone (9p TelVysion probe #05J03-009) mapping to chromosome 9 was labeled in green using Spectrum Green (Vysis, Abbott Molecular, Des Plaines, IL). Both probes were hybridized to 46,XY control metaphases. The slides were denatured with 50% formaldehyde/2xSSC at 80°C for 1.5 minutes and then dehydrated with serial ethanol washing steps (ice cold 70, 90, and 100% for 3 minutes each). The probes were denaturated in the hybridization solution (50% dextran sulfate/2xSSC) at 85°C for 5 minutes, applied to the slides, and incubated overnight at 37°C in a humidified chamber. The slides were then washed 3 times for 5 minutes with 50% formamide/2X SSC, 1X SSC and 4xSSC/0.1%Tween. Slides were dehydrated with serial ethanol washing steps (see above) and mounted with ProLong Gold antifade reagent with DAPI (Invitrogen, Carlsbad, CA) for imaging. Image acquisition is described in **Supplemental Experimental Procedures**.

### RNA:DNA hybrid immunoprecipitation (RDIP)

The cell culture conditions for IMR-90 and HEK 293T cells are described in the **Supplemental Experimental Procedures**. Whole cell nucleic acid was isolated from HEK 293T cells and IMR-90 cells through a modified salting out extraction protocol [85]. Nucleic acid was sonicated to an average size of 400-600 bp using the Covaris sonicator. The fragmented nucleic acid was then treated with RNase I (Ambion AM2294) to remove any ssRNA from the sample, phenol/chloroform purified and re-suspended in EB buffer. Part of the nucleic acid sample was set aside as an untreated input sample for comparative sequencing. Three micrograms of nucleic acid sample was then incubated overnight with the S9.6 antibody, following which the RNA:DNA hybrids were enriched by immunomagnetic precipitation using Dynabeads (M-280 Sheep anti-mouse IgG). The sample was then extracted through phenol/chloroform purification, precipitated in the presence of glycogen and re-suspended in EB buffer. A complete detailed protocol is available in the **Supplemental Experimental Procedures**. Enrichment of predicted peaks in the RDIP product was validated using quantitative PCR (Quanta PerfeCTa SYBR Green Fastmix). The primer sequences used are provided in **Table S3**.

### Directional RDIP-seq

Using RDIP and input material, directional RDIP-seq libraries were prepared using elements of a directional RNA-seq protocol modified from a previously published approach [50]. Starting the library preparation at the second strand synthesis step, the RNA of the RNA:DNA hybrid was nicked using RNase H treatment to serve as a primer for the DNA polymerase. The second strand was formed while incorporating dUTP to allow for directional sequencing and the identification of the RNA strand of the RNA:DNA hybrid. Next, the ends of fragments were repaired, adenosine tails added, and Illumina Tru-Seq strand-specific adaptors ligated (adaptor sequences in **Supplemental Table S4**). UNG treatment was utilized to degrade the dUTP-containing RNA strand of the RNA:DNA hybrid, and barcoded PCR primers were used to amplify the library while maintaining directionality. The complete RDIP-seq protocol is available in the **Supplemental Experimental Procedures**.

Prior to sequencing, the libraries were analyzed for quality of preparation using an Agilent Bioanalyzer high-sensitivity chip. Libraries were multiplexed and combined for sequencing using Illumina HiSeq 2500 150 bp paired-end sequencing in our institutional Epigenomics Shared Facility. Fastq files were generated through the Illumina CASAVA pipeline (v1.8). Sequencing reads were then run through the Wasp System (WASP v3.1.5 rev. 6632) hosted pipeline for primary data processing, as follows. The reads were aligned to the hg19 reference genome using Bowtie (v0.12.7), using non-default parameters of --tryhard (increasing the number of attempts bowtie uses to find an alignment and number of backtracks), -I 50 (the minimum insert size in basepairs for valid paired-end alignments) and -X 650 (the maximum insert size for valid paired-end alignments). Alignments were generated in SAM format, which were then transformed into BAM files using Samtools (version 0.1.8). The aligned sequences in BAM format had PCR duplicates removed, and peaks were called based on input and IP files using MACS v1.4.2 [86]. RDIP-seq peaks for IMR-90 cells and two datasets for HEK 293T cells were then analyzed using the program CHANCE for quality of immunoprecipitation [87]. Based on the results of CHANCE, we discarded one of the HEK 293T datasets and continued on with one set of peaks for each cell line. All peaks containing “N” nucleotides were discarded. Custom code and parameters for this analysis can be found on our GitHub resource in the file “Peak Calling.” Motif analysis of RNA:DNA hybrid peaks is described in the **Supplemental Experimental Procedures**.

### R-loop validation through non-denaturing bisulphite conversion

RDIP-seq peaks were validated through non-denaturing bisulphite conversion. Whole cell nucleic acid was isolated from HEK 293T cells through a modified salting out extraction protocol as outlined in the **Supplemental Experimental Procedures**. Nucleic acid was digested with EcoRV-HF. Non-denaturing bisulphite treatment was performed according to a previously published protocol [51]. Regions of interest were amplified through PCR after denaturing or non-denaturing bisulphite treatment using primers to converted or unconverted DNA. The PCR product was purified, cloned using a TOPO-TA cloning kit (Life Technologies) and sequenced. The primer sequences used in non-denaturing bisulphite validation for this study are provided in **Table S5**.

### Directional RDIP-seq strandedness analysis

Due to using directional sequencing through the incorporation of dUTP, we were able to determine the RNA-derived sequence of the RNA:DNA hybrids. To do this, we used the BAM flag information describing our aligned sequences (http://broadinstitute.github.io/picard/). The second read in the pair, representing the sequence derived from the RNA strand following degradation using UNG of the dUTP-incorporated complementary strand, has the bit flag identifiers of 163 or 147, indicating that it maps to the top or bottom strand of the reference genome, respectively. By measuring the number of RNA reads aligned to the top or bottom reference strand for each peak, we could assign each RDIP-seq peak a “strandedness” value, with +1 being all RNA-derived reads aligned to the top strand and -1 all RNA-derived reads aligned to the bottom strand. We removed the small minority (10%) of peaks with intermediate values of strandedness to decrease what we presumed to be experimental noise in our data set. Custom code for this analysis can be found on our GitHub resource in the file “Determining RNA Strand and Minus10 files.”

### RNA-seq of HEK 293T cells and IMR-90 cells

RNA was isolated from HEK 293T and IMR-90 cells using TRIzol extraction. Four biological replicates from each cell line were DNase treated, and Ribo-Zero rRNA removal (Ribo-Zero, Epicentre) was utilized for three of the four RNA samples, leaving a non-Ribo-Zero depleted sample for rRNA expression analysis. RNA-seq libraries were prepared using a directional RNA-seq protocol modified from a prior published approach [50] and detailed in the **Supplemental Experimental Procedures** Directional Whole Transcriptome Sequencing protocol. Prior to sequencing, the libraries were assessed for quality using an Agilent Bioanalyzer high-sensitivity chip. The samples were multiplexed and sequenced using 100 bp single-end read sequencing on the Illumina HiSeq 2500 in our institutional Epigenomics Shared Facility. The TruSeq adaptor sequences used in this assay are provided in **Supplemental Table S6**.

After sequencing, fastq file generation was completed using the Illumina CASAVA pipeline (v1.8). Post-sequencing analysis was performed using the WASP pipeline (v3.1.5 rev. 6632), involving read alignment using gsnap (2012-07-20), with htseq (v0.5.3p3) used to determine read quantitation. Biological replicates were normalized using DESeq (Bioconductor) and RefSeq gene identifiers were assigned using biomaRt. Only gene expression assigned a RefSeq identifier was used for further analysis. Custom code for this analysis can be found on our GitHub resource under the file “RNAseq Analysis.”

### Ribosomal DNA analysis

In order to align our RDIP-seq reads to the rDNA repeating unit, we used the alignment approach of Zentner and colleagues [1]. We added the rDNA repeating unit fasta file (gi|555853|gb|U 13369.1|HSU13369) to the start of the hg19 chromosome 13, replacing the telomeric “N” nucleotides. Duplicate reads were removed from the IMR-90 RDIP-seq and input fastq files using a custom perl script provided by Zentner and colleagues [1], and the remaining reads were aligned to the hg19+rDNA genome file using Bowtie. Wiggle tracks were then created using FSeq, and counts representative of the reads aligned to the rDNA portion of chromosome 13 were isolated. The RDIP-seq wiggle track values were normalized by subtracting the input values from the RDIP values. The same pipeline was used to align the IMR-90 RNA-seq samples that did not have prior Ribo-Zero depletion to the rDNA sequence. Processed histone mark datasets from K562 cells for rDNA were provided by Zentner and colleagues [1], and averaged across 50 bp windows across the rDNA repeating unit. Custom code for this analysis can be found on our GitHub resource under the file “Figure 2 - rDNA figure with Zentner histone marks” and the custom perl script under “Zentner removeDupsFromFastQ Perl Script.”

### Regression models of RNA:DNA hybrid peak density

We used LASSO regularized linear regression to explore the relationship between the density of RNA:DNA hybrid peaks in 500 kb windows and genomic features associated with transcription and regulation. LASSO regression fits a linear model subject to a constraint on the sum of the regression coefficients [66]. The LARS algorithm, implemented in the *LARS* R package, was applied to determine the Lasso path. This algorithm provides the optimal values of the regression coefficients as the constraint on the sum of the coefficients is progressively relaxed [67]. Tight constraint on the sum of the coefficients enforces sparseness on the model with the number of covariates in the model increasing as this constraint is relaxed. The covariance test statistic [68], implemented in the *covTest* R package, was used to test the significance of each additional covariate when it enters the model.

### Co-Immunoprecipitation of RNA:DNA Hybrid Binding Proteins (CoIP)

Native chromatin was isolated using a sucrose gradient from HEK 293T cells. Chromatin was incubated overnight with S9.6 antibody or a non-specific control antibody (β-actin, Sigma A5441), following which immunoprecipitation was performed on each sample using immunomagnetic precipitation (Dynabeads M-280 Sheep anti-mouse IgG). RNA:DNA hybrid-binding protein complexes were then eluted using RNA:DNA hybrid oligonucleotides, with DNA:DNA oligonucleotides as a control. The oligonucleotide sequences used in this assay are provided in **Supplemental Table S7**. The resulting enriched proteins were run on a 12% polyacrylamide gel, stained with GelCode Blue (Life Technologies 24594) and tested using Mass Spectrometry (MS). Proteins which were considered to bind specifically to RNA:DNA hybrids were defined as those only present in the S9.6 immunoprecipitated sample and eluted with the RNA:DNA oligonucleotides, removing any proteins also present in the control samples (those isolated with the β-actin antibody, and with the S9.6 antibody eluted with the DNA:DNA oligonucleotides). This analysis was performed using Scaffold3 proteome software [88]. Peptide counts were assigned to each protein identified through mass spectrometry by measuring the quantity of the identified peptides by their spectra, and filtered by those peptides that also occurred in negative control experimental samples. Candidate proteins identified by mass spectrometry were then validated using Western blotting using the antibodies described in **Supplementary Table S8**.

### Custom Code

Analysis of RDIP-seq, RNA-seq, and code for all figures are included and annotated at: https://github.com/GreallyLab/Nadel-et-al.-2015

## DATA ACCEES

The data generated are all available through the Gene Expression Omnibus, accession number GSE68953 (http://www.ncbi.nlm.nih.gov/geo/query/acc.cgi?acc=GSE68953).

## ABBREVIATIONS

BAC bacterial artificial chromosome; ChIP chromatin immunoprecipitation; CpG cytosine-guanine dinucleotide; DAPI 4',6-diamidino-2-phenylindole; DNA deoxyribonucleic acid; DRIP DNA:RNA immunoprecipitation; dsDNA double stranded DNA; dUTP deoxyuridine triphosphate; EMSA electrophoretic mobility shift assay; FISH fluorescence *in situ* hybridization; H3K27me3 histone H3 lysine 27 trimethylation; H3K9me3 histone H3 lysine 9 trimethylation; H3S10 histone H3 serine 10; IGS intergenic spacer of ribosomal DNA repeat unit; LARS least angle regression; LASSO least absolute shrinkage and selection operator; LINE long interspersed nuclear element; lncRNA long non-coding RNA; mRNA messenger RNA; NOR nucleolar organising region; PCR polymerase chain reaction; polyA polyadenylation; RDIP RNA:DNA hybrid immunoprecipitation; rDNA ribosomal DNA; RNA ribonucleic acid; ssDNA single stranded DNA; TERRA telomeric repeat containing RNA; TF transcription factor; UNG uracil N-glycosylase.

## AUTHOR CONTRIBUTIONS

J.M.G. and J.J. designed the original project. J.M.G., R.A. and J.N. designed the original experiments and the analytical approaches. J.N. executed the experiments and analyzed results. J.M.G. and J.N. wrote the manuscript. C.L., N.A.W. and Z.Z. contributed to analysis. P.O. and A.G. performed the motif analysis. H.S. performed the immunofluorescence studies, and the group of C.M. performed the FISH studies. C.S. performed the LASSO analysis. All authors read and approved the final manuscript.

## ADDITIONAL DATA FILES

Two additional data files are provided, one with supplementary data and methods, the other with supplementary tables.

## AUTHOR INFORMATION

Rodoniki Athanasiadou’s current address: New York University, Center for Genomics and Systems Biology, Department of Biology, 12 Waverly Place, New York, NY 10003, USA

Christophe Lemetre’s current address: Integrated Genomics Operation, Memorial Sloan-Kettering Cancer Center, New York, NY 10065, USA.

## ACKNOWLEDGEMENTS

The project was funded by NIH R21 GM101880 grant to JMG. Einstein core facilities involved were the Genome Imaging Core, the High-Performance Computing Core, the Epigenomics Shared Facility, the Proteomics Core Facility, the Monoclonal Antibody Core Facility and the Genomics Core Facility, with support from the Albert Einstein Cancer Center (P30CA013330) and the Center for Epigenomics. JN was supported by the Training Program in Cellular and Molecular Biology and Genetics, (NIH T32 GM007491) and NAW by the Medical Student Training Program (NIH T32 GM007288). We thank Drs. Gabe Zentner and Peter Scacheri at Case Western Reserve University for sharing code and data to allow us to perform the rDNA studies as comparably as possible with their prior work.

